# Epigenetic anti-cancer treatment with a stabilized carbocyclic Decitabine analogue

**DOI:** 10.1101/2022.01.19.476908

**Authors:** Franziska R. Traube, Natércia F. Brás, Wynand P. Roos, Corinna C. Sommermann, Tamara Diehl, Robert J. Mayer, Armin R. Ofial, Markus Müller, Hendrik Zipse, Thomas Carell

## Abstract

5-Aza-2’-deoxycytidine (Decitabine, AzadC) is a nucleoside analogue, which is in clinical use to treat patients with myelodysplastic syndrome or acute myeloid leukemia. Its mode of action is unusual because the compound is one of the few drugs that act at the epigenetic level of the genetic code. AzadC is incorporated as an antimetabolite into the genome and creates covalent, inhibitory links to DNA methyltransferases (DNMTs) that methylate 2’-deoxycytidine (dC) to 5-methyl-dC (mdC). Consequently, AzadC treatment leads to a global loss of mdC, which presumably results in a reactivation of silenced genes, among them tumor suppressor and DNA damage response genes. Because AzadC suffers from severe instability, which limits its use in the clinic, a more sophisticated AzadC derivative would be highly valuable. Here, we report that a recently developed carbocyclic AzadC analogue (cAzadC) blocks DNMT1 in the AML cell line MOLM-13 as efficient as AzadC. Moreover, cAzadC has a surprisingly strong anti-proliferative effect and leads to a significantly higher number of double strand breaks compared to AzadC, while showing less off-target toxicity. These results show that cAzadC triggers more deleterious repair and apoptotic pathways in cancer cells than AzadC, which makes cAzadC a promising next generation epigenetic drug.

---

Acute myeloid leukemia (AML) is a severe highly diverse cancer of the blood building system and very difficult to treat.^[1]^ AzadC **1** (Fig. 1) is an orphan drug used to treat AML in older, medically non-fit patients with a significant beneficial effect on patient outcome.^[2]^ The broader use of AzadC **1** is currently limited due to the high hematotoxicity of the compound, which sets tight boundaries to the dosing.^[3]^ The treatment requires repeated administration cycles, which however, can go on for several years.^[2, 4]^ AzadC **1** operates at the epigenetic level and has potential also for the treatment of cancers other than AML.^[5–6]^ The compound is converted in cells into the corresponding triphosphate followed by a DNA-polymerase-based integration into the genome of dividing cells.^[7–9]^ After genomic incorporation, AzadC **1** represents an abnormal base, which is removed together with its degradation products by the base excision repair (BER) system.^[10]^ BER requires a family of glycosylases, which attack the anomeric C1’ position with a nucleophile to cleave the glycosidic bond. This ultimately results in a DNA single-strand break (SSB), which is quickly repaired.^[11]^ In addition, genomically incorporated AzadC **1** inhibits DNA methyltransferases (DNMTs).^[7]^ These enzymes initially catalyze a nucleophilic attack at the C6 position of dC, which gives a covalent intermediate that is subsequently methylated at C5 by the cofactor SAM. In the last step, the DNMT enzyme is released.^[12]^ This DNA methylation in promoter regions causes a permanent silencing of the corresponding genes.^[13–14]^ Due to the exchange of the −CH at the C5 position to N, AzadC **1** leads to covalent trapping of the DNMTs because the protein cannot be released anymore after the methylation reaction.^[7]^ This trapping creates DNA-protein crosslinks, which results in a reduction of the global mdC levels.^[9]^ AzadC **1** has therefore three levels of action. First, it triggers BER-mediated repair of the abnormal base itself or of its degradation products. Second it leads to DNA-protein adducts, which are repaired by Fanconi anemia-dependent homologous recombination (FA pathway).^[10, 15]^ Finally, it operates at the epigenetic level by causing global DNA demethylation.

**Figure 1.**
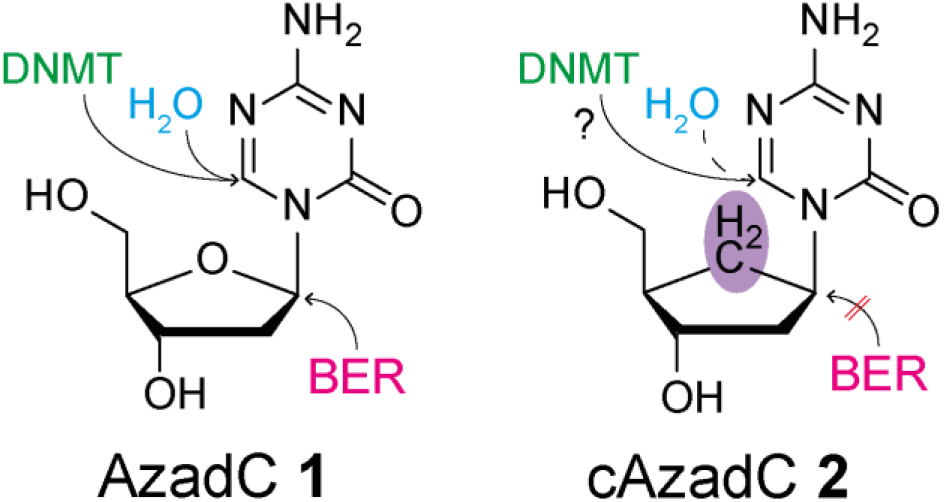
Presentation of 5-aza-2’-deoxycytidine (AzadC **1**) and a carbocyclic derivative thereof (cAzadC **2**). AzadC **1** is already in use in the clinic as an orphan drug against different forms of leukemia. For AzadC **1**, nucleophilic attacks occur at position 6 of the nucleobase by water or DNMT enzymes or at the anomeric C1’-center by a BER glycosylase. For cAzadC **2**, hydrolysis is dramatically slowed down (H_2_O dashed line) and the nucleophilic attack at position C1’ by glycosylases is not possible anymore.

The problem with AzadC **1** is its inherent instability against hydrolytic deactivation especially at basic or acidic pH, which is often a hallmark of the microenvironment of solid tumors.^[16–17]^ We believe that stabilized 5-azacytosine based nucleosides with a more defined drug profile would have a chance to become broadly beneficial as anti-cancer agents. Recently, we reported in this direction a carbocyclic version of AzadC (cAzadC **2**), in which the ribose O-atom is replaced by a CH_2_-group (Fig. 1). We observed that cAzadC **2** is substantially more stable than AzadC **1** towards hydrolysis.^[17]^ However, in a mouse embryonic stem cell model, cAzadC **2** required higher dosage and longer incubation times to show a demethylation effect comparable to AzadC **1** raising the question whether cAzadC **2** can inhibit the DNMTs as efficiently as AzadC **1** in cancer cells.

First, we determined the rate constant of a nucleophilic attack of a hydroxide ion at position C6. In principle, we observed the same reaction kinetics for AzadC **1** (Fig. S1a) and cAzadC **2** (Fig. S1b), but in accordance with our previous study, the rate constant k was 59 times higher for AzadC **1** (k = 2.13 × 10^−2^ M^−1^ s^−1^) compared to cAzadC **2** (k = 3.6 × 10^−4^ M^−1^ s^−1^). To obtain quantitative data on the potentially reduced electrophilic activity of the C6 position in **2**, we modelled the corresponding reaction products after the nucleophilic attack of the hydroxyl group on C6 from the representative optimized geometries of AzadC **1**, cAzadC **2** and dC **3**. After conformational analysis of **1**, **2** and **3** in an aqueous environment (see Schemes S1 – S2, Tables S1 – S4 and Fig. S2 for details), the respective product geometries were optimized at the SMD(water)/B3LYP-D3/6-31+G(d,p) level of theory. Fig. 2 illustrates the most thermodynamically stable conformers for each of the three systems together with their respective conformational free energies. The reaction energies ΔG_R_, which are shown for the best conformers, were calculated relative to the best conformer of the neutral parent plus free hydroxide in water. Conformations with intramolecular hydrogen bonds between the C5’-OH and the O2 of the base (*syn* conformation) were the most favorable for all three systems studied here. For both AzadC **1** and dC **3**, we also found conformers with an intramolecular H-bond between the added nucleophile hydroxyl group and the ring oxygen atom of ribose, which is absent in cAzadC **2**, resulting in an energetically favored six-membered ring. The reaction energies for hydroxide addition to the C6 carbon atom are most favorable for AzadC **1** (ΔG_R_ = −0.1 kJ mol^-1^), followed by cAzadC **2** (ΔG_R_ = +10.4 kJ mol^-1^), and finally dC **3** (ΔG_R_ = +55.8 kJ mol^-1^). The reaction energy difference between AzadC **1** and cAzadC **2** of ΔΔG_R_ = 10.5 kJ mol^-1^ is remarkably similar to the activation free energy difference observed between those two systems in water at 20°C of ΔΔG_act_ = 9.9 kJ mol^-1^. We therefore concluded that the reactivity difference between AzadC **1** and cAzadC **2** towards hydrolysis presumably results from the stability differences between the initially formed hydroxide adducts at the C6 position (**1**@H_2_O versus **2**@H_2_O).

**Figure 2.**
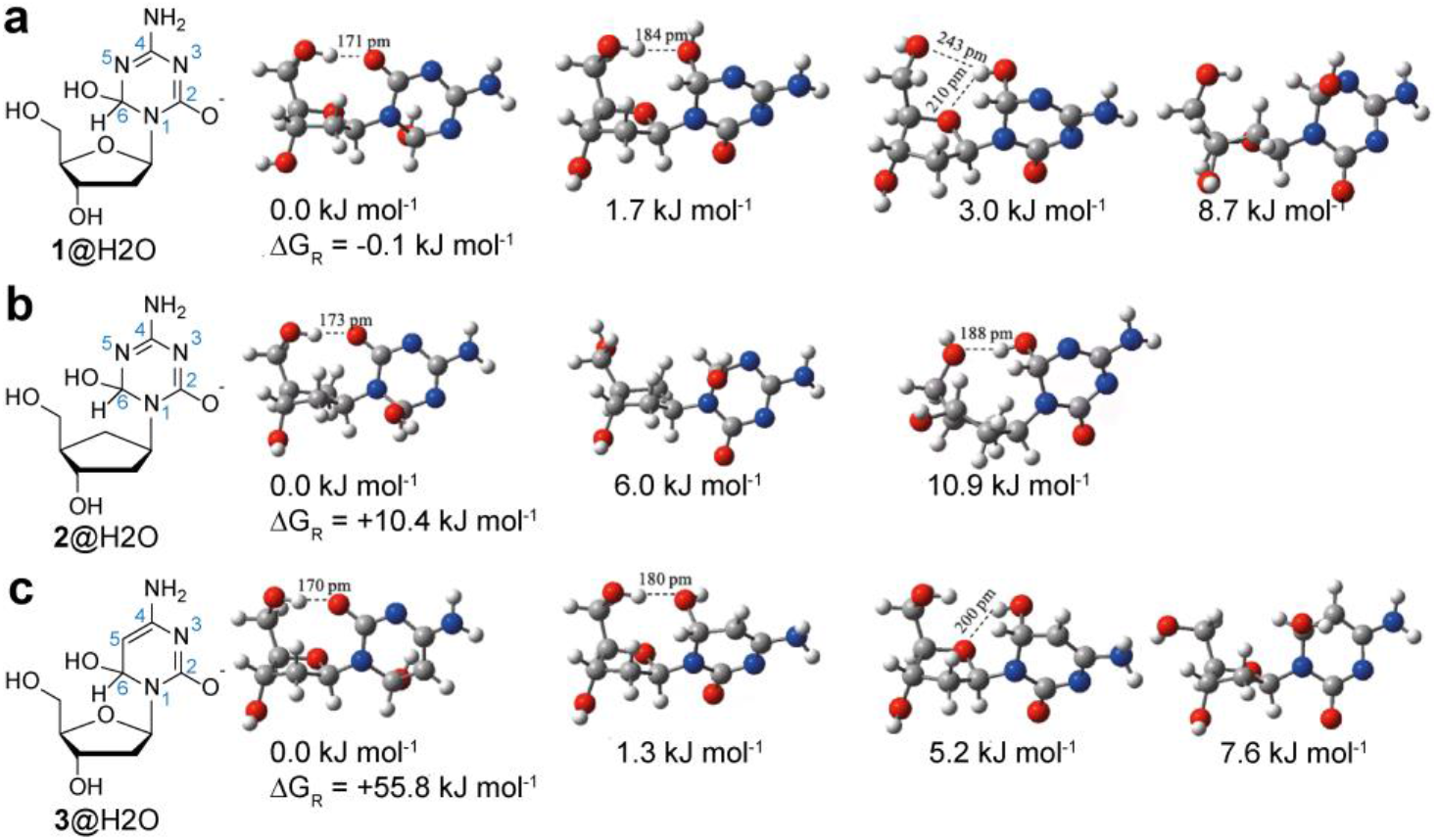
Thermodynamically most favored geometries of AzadC **1**@H_2_O (**a**), cAzadC **2**@H_2_O (**b**) and dC **3@**H_2_O (**c**) obtained at the SMD(water)/B3LYP-D3/6-31+G(d,p) level of theory. The Gibbs energy of each molecule and the reaction energies are displayed.

Next, we aimed to assess how AzadC **1** and cAzadC **2** inhibit DNMTs in cultured cells. Inhibition of DNMTs by crosslinking them to the DNA is followed by crosslink removal as part of the repair process and subsequent DNMT degradation (Fig. 3a).^[18–19]^ Therefore, efficient DNMT inhibition by 5-azacytidines results in reduced amounts of DNMT enzymes, which can be detected by western blotting. To test the ability of AzadC **1** and cAzadC **2** to deplete the DNMT pool, we decided to use the AML cell line MOLM-13 as a model system, since AzadC **1** is primarily applied in the clinic against AML and myelodysplastic syndrome. We treated the MOLM-13 for 40 h with 0.5 μM AzadC **1** or 0.5 μM cAzadC **2**. MOLM-13 that were treated only with 0.03% DMSO served as a control. Afterwards, the cells were lysed and the nuclear proteins were isolated as previously described. We then analyzed the amount of DNMT1, which is the most abundant DNMT in MOLM-13 by western blotting.^[20–21]^ For both compounds, we detected a clear depletion of DNMT1 and to our surprise, the effect was stronger for cAzadC **2** compared to AzadC **1** (Fig. 3b, Fig. S3). This shows that the higher hydrolytic stability of cAzadC **2** does not affect its reactivity towards DNMTs *in cellulo*. Next, we investigated how application of increasing concentrations of AzadC **1** or cAzadC **2** (0.5 μM, 1.0 μM or 3.0 μM) for 72 h influence the global mdC levels in MOLM-13 using our previously reported UHPLC-QQQ-MS method (Fig. 3c).^[22]^ We observed the expected substantial reduction of mdC for all concentrations and both compounds. At higher concentrations (3.0 μM), AzadC **1** seems to show a slightly better reduction (−57%) compared to cAzadC **2** (−40%), but the difference was not statistically significant considering the overall reduction compared to the untreated control.

**Figure 3.**
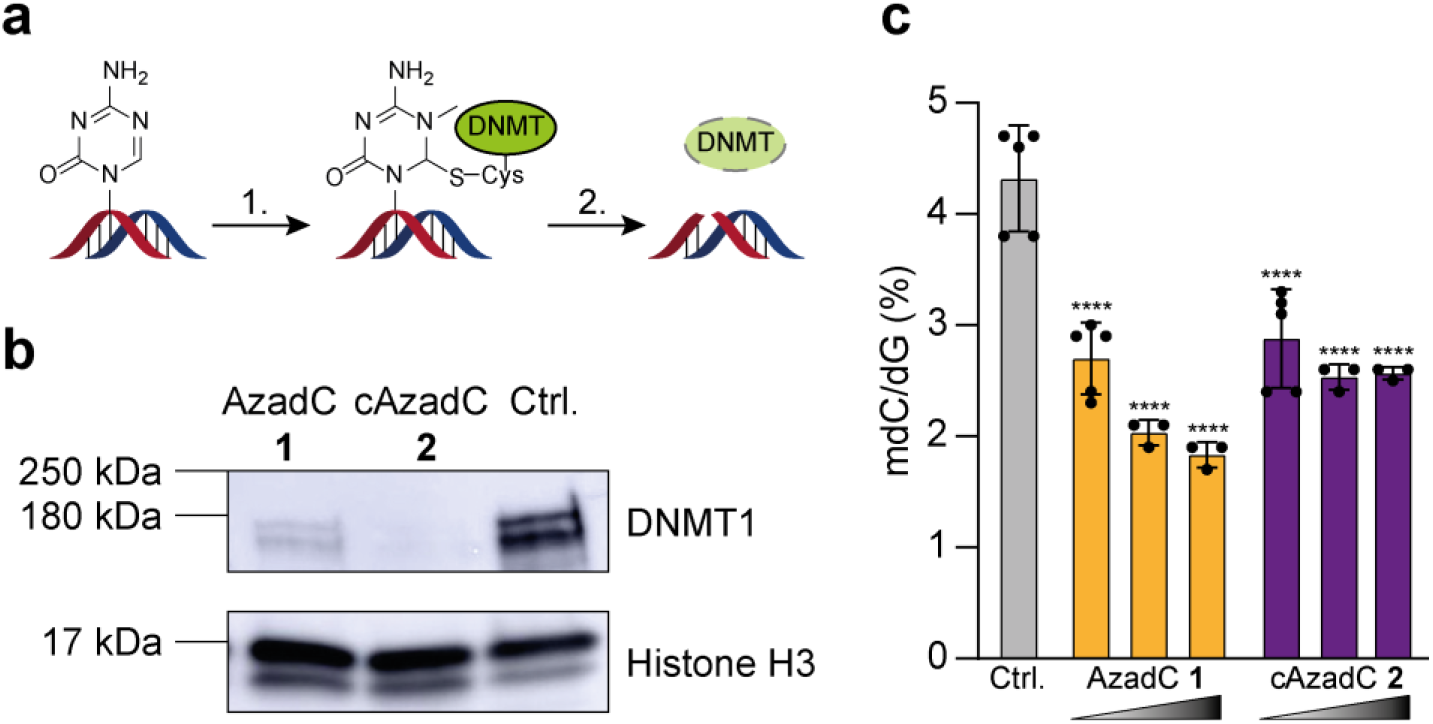
Inhibition of DNMT enzymes in MOLM-13 by AzadC **1** or cAzadC **2**. **a** Formation of DNA-protein crosslinks between DNMT enzymes and 5-aza-cytosines (1.) is followed by removal of the crosslink and DNMT degradation (2.). **b** Western blot against DNMT1 and histone H3 as a loading control. 15 μg of nuclear lysate of MOLM-13 that were treated for 40 h with 0.5 μM of AzadC **1** or cAzadC **2** were used. Untreated cells served as a negative control. C Global mdC levels determined by UHPLC-QQQ-MS of MOLM-13 that were treated for 72 h with either 0.5 μM, 1.0 μM or 3.0 μM of AzadC **1** or cAzadC **2** (increasing concentrations are represented by the triangle). Ordinary one-way ANOVA combined with Tukey’s multiple comparisons test was used to analyze differences in the mdC levels between the different conditions. The adjusted p-value of the treated cells compared to the Ctrl. is displayed, **** p_adj_-value < 0.0001. Each dot represents measurements from one biologically independent experiment.

To compare AzadC **1** and cAzadC **2** in their ability to trigger cell death in the MOLM-13, we measured the viability of MOLM-13 with increasing exposure time (24 – 72 h) and concentrations (0.5 – 3.0 μM) of AzadC **1** and cAzadC **2** using a methylthiazolyldiphenyl-tetrazolium bromide (MTT) based metabolic activity assay.^[23]^ The data show that both compounds lead to strongly reduced metabolic activity indicating advanced cell death with increased exposure time at all concentrations that were examined (Fig. 4a). In this assay the stabilized compound cAzadC **2** showed a stronger effect than AzadC **1**, although the difference between AzadC **1** and cAzadC **2** was not statistically significant when considering the overall reduction of metabolic activity compared to the untreated control. After 72 h of 3.0 μM cAzadC **2** exposure, the metabolic activity of the MOLM-13 was reduced to less than 10% for cAzadC **2**, while still 20% residual metabolic activity was observed for MOLM-13 treated with 3.0 μM of AzadC **1**. Given that both compounds have a dual function consisting of mdC reduction, which may reactivate silenced tumor suppressor genes, and formation of DNA-protein crosslinks, which is a severe form of DNA damage, the better performance of cAzadC **2** compared to AzadC **1** on the AML cell line might be caused by different repair profiles. All types of DNA repair involve SSBs as intermediates that are quickly repaired by the different repair machineries to avoid formation of the more deleterious double-strand breaks (DSBs), a process which is dramatically accelerated by the accumulation of SSBs.^[24]^ The exchange of the ribose O in AzadC **1** to CH_2_ in cAzadC **2** has the effect that repair glycosylases of BER cannot cleave the glycosidic bond anymore to create an abasic site. To investigate the level of DNA damage in MOLM-13 caused by 0.5 μM of AzadC **1** or cAzadC **2** exposure after 40 h, we performed first a western blot against γH2AX, which is a histone variant, phosphorylated at position Ser-139, that is placed a sites of DSBs to recruit the repair machinery. γH2AX therefore represents an early marker of severe DNA damage (Fig. 4b, Fig. S3).^[25]^ As expected, the western blot shows highly increased levels of γH2AX, confirming that both compounds induced substantial DNA damage after 40 h. We then quantified the extent of SSBs and DSBs in individual cells after longer exposure times (60 h). To this end we performed a single-cell gel electrophoresis comet assay.^[26–27]^ For the assay, MOLM-13 cells were incubated for 60 h with either 0.5 μM of AzadC **1** or cAzadC **2**. Untreated cells served as a negative control, while cells treated with 150 μM of t-BuOOH for 1 h served as a positive control (Fig. S4a, b). Afterwards, the cells were embedded in agarose placed on a slide and exposed to a lysis buffer. The DNA was subsequently denatured and subjected to electrophoresis followed by fluorescence microscopy, where it creates an image like a comet. The head of the comet is composed out of intact DNA, while the tail is generated by broken DNA. The longer the tail, the more broken DNA was generated by the added substance. Importantly, when the assay is conducted under neutral conditions, the tail is predominantly caused by DSBs, while under alkaline conditions SSBs and DSBs can be observed in theory. However, the contribution of the DSBs to the tail length is only marginal compared to the SSBs under alkaline conditions. The comet assay consequently allows to measure which types of DNA strand breaks are generated by a given compound. The data obtained for AzadC **1** and cAzadC **2** are depicted in Fig. 4c and d. For both compounds, the levels of SSBs were increased relative to the untreated control (Fig. 4c), but the increase was only significant for cAzadC **2** when comparing the tail intensities among all three treatments (untreated, AzadC **1** and cAzadC **2**). Furthermore, we detected for cAzadC **2** highly increased levels of the deleterious DSBs not only compared to the untreated control but also compared to treatment with equal amounts of AzadC **1** (Fig. 4d). The combined comet assay data suggest that AzadC 1 treatment either leads to less lesions than cAzadC **2** treatment or that lesions caused by AzadC **1** can be repaired more efficiently by the MOLM-13. Although the glycosidic bond of cAzadC **2** cannot be cleaved by BER glycosylases and hydrolytic degradation is dramatically reduced, which prevents the formation of abasic sites, treatment with cAzadC **2** lead to significantly more damage than treatment with AzadC **1**. Existing literature suggests that abasic sites caused by AzadC 1 are quickly repaired when the BER machinery is intact.^[10]^ This can explain the comet assay results. Important is the observation that the levels of DSBs after 60 h were strongly increased after cAzadC **2** treatment. This shows that cAzadC **2** causes DNA lesions that remain unrepaired to a larger extent and are therefore highly deleterious for the cells.

**Figure 4.**
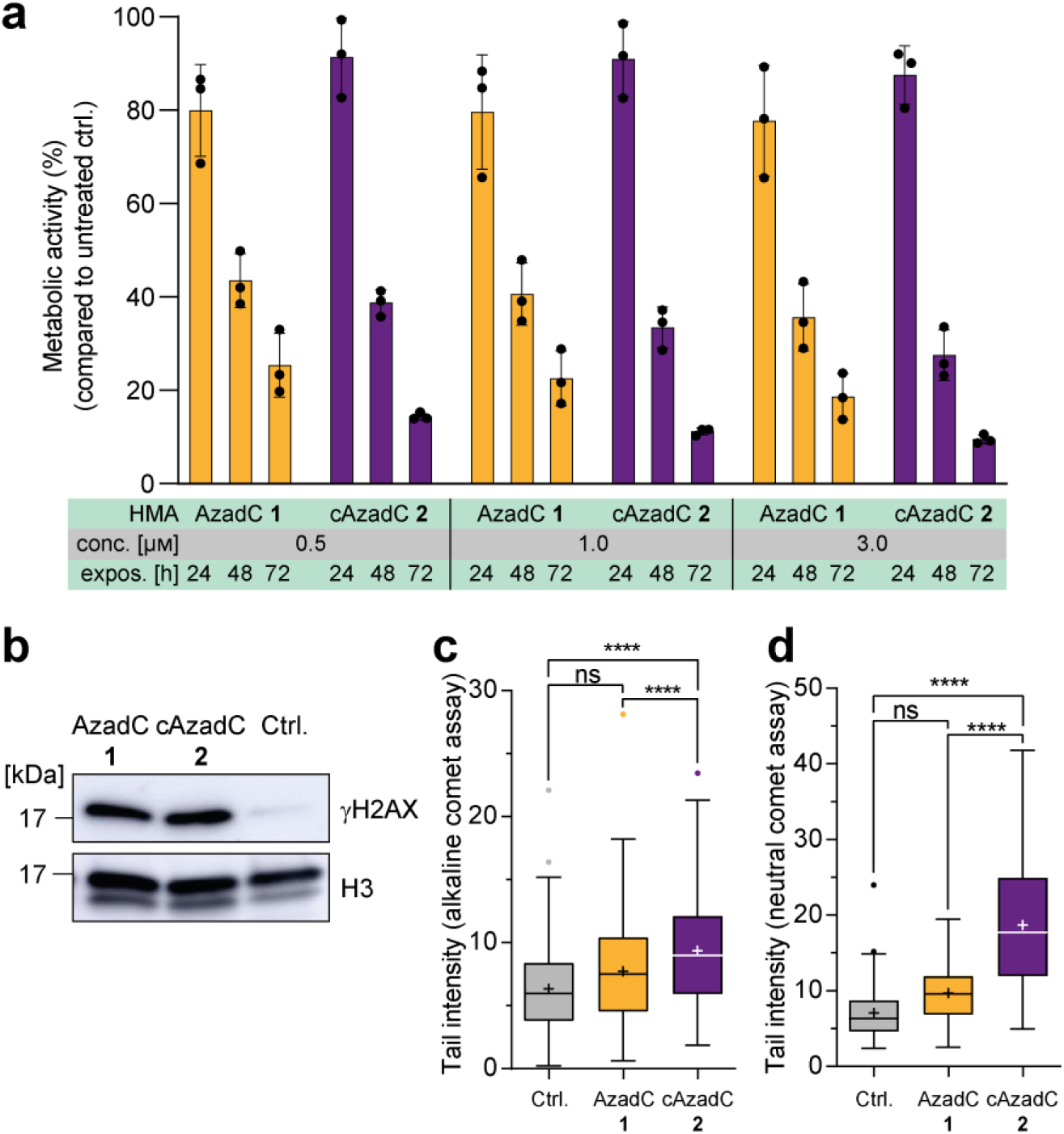
Cellular viability and DNA damage in MOLM-13 after AzadC **1** or cAzadC **2** treatment. **a** MTT assay to check for metabolic activity as an indicator for cell viability and proliferation of MOLM-13. The cells were treated for 24 h, 48 h or 72 h with 0.5 μM, 1.0 μM or 3.0 μM of either AzadC **1** or cAzadC **2** before the assay was performed. Untreated cells served for each measurement as a reference to normalize the metabolic activity of the treated cells. Each dot represents one biologically independent sample. **b** Western blot against γH2AX and histone H3 as a loading control. 15 μg of nuclear lysate of MOLM-13 that were treated for 40 h with 0.5 μM of AzadC **1** or cAzadC **2** were used. Untreated cells served as a negative control. **c** and **d** MOLM-13 were treated for 60 h with either 0.5 μM of AzadC **1** or cAzadC **2** before the comet assay was performed. **c** Result of the alkaline comet assay to determine DNA single-strand breaks. d Result of the neutral comet assay to determine the DNA double-strand breaks. **c** and **d** Brown-Forsythe and Welch ANOVA tests combined with Dunnett’s T3 multiple comparisons test were used for the analysis. ns p_adj_-value ≥ 0.5, **** p_adj_-value < 0.0001. Three biologically independent experiments were performed for each condition and 50 tail measurements were performed within each biological replicate.

In summary, our decitabine-derivative cAzadC **2** is not only chemically significantly more stable than AzadC **1**, which makes administration easier, but it also shows increased activity to induce cell death of cancer cells despite the higher stability. Initial data about potential off-target toxicity of cAzadC **2** seem to indicate that the compound can be dosed much higher than AzadC **1** (Fig. S5). This, together with the here reported data suggest that cAzadC **2** could open a new chapter in our desire to establish epigenetic anticancer treatments.

## Acknowledgements

We thank the Deutsche Forschungsgemeinschaft (DFG) for financial support via CRC1309 (Project ID 325871075) and CRC1361 (Project ID 893547839). This project has received funding from the European Research Council (ERC) under the European Union’s Horizon 2020 research and innovation program under grant agreement No 741912 (EpiR). We are grateful for additional funding from the Volkswagen Foundation. N. F. Brás thanks Portuguese Foundation for Science and Technology for a research grant (CEECIND/02017/2018). We thank Luis de la Osa de la Rosa for initial assistance with the MTT assays and passaging of MOLM-13.

