## Supplementary Information for "Epigenetic anti-cancer treatment with a stabilized carbocyclic Decitabine analogue"

### Materials and Methods

Comparison of the reactivity of AzadC **1** and cAzadC **2**

The reactivity of AzadC **1** and cAzadC **2** was determined using stopped-flow spectrophotometry using the absorbance changes in the UV region. For both compounds, the attack of hydroxide ions was monitored in aqueous solution at 20 °C (pH 11 – 12). To simplify the evaluation, the kinetic measurements were performed under 1st order conditions, i.e., the hydroxide concentrations were chosen to be high enough to be considered constant during the course of reactions (12.5 mᴍ KOH and 0.35 mᴍ AzadC **1** or 5.0 mᴍ KOH and 0.41 mᴍ cAzadC **2**).

The UV spectra in Fig. S1a show a two-step reaction between AzadC 1 and $OH^-$. In the first step of the reaction sequence, a new species was formed within a few seconds with a clearly pronounced absorption at 248 nm (left panel, brown to green UV spectra). In a slower subsequent reaction, the peak at 248 nm was disappearing while a new species with an absorption maximum at 225 nm was formed (left panel, green to black UV spectra. The isosbestic point (UV spectra green to black) indicates that the decay of the AzadC/$OH^-$ primary adduct to the secondary product proceeded without forming any intermediates. The first-order rate constant $k_{obs}$ for the temporal change of the absorption at 248 nm by formation of the primary adduct was determined at different $OH^-$ concentrations. The rate for the formation of the primary adduct showed a clear linear dependence on the concentration of the nucleophilic $OH^-$ ions. From the slope of the plot (Fig. S1a, right panel), a rate constant $k = 2.13 \times 10^{-2}$ ᴍ$^{-1}$ $s^{-1}$ could be determined for the attack of the $OH^-$ ions on AzadC.

In principle, the observed kinetics for the reactions of the $OH^-$ ions with cAzadC **2** are analog to those for $OH^-$/AzadC (Fig. S1b). There is also an initial increase of absorption at 248 nm (left panel, brown to green UV spectra), which is caused by the formation of the primary adduct. However, as the reaction continues, the adduct disappears, leaving behind a second species with an absorption maximum at 228 nm (left panel, green to black UV spectra). Since the formation of the primary adduct was much slower in this case, but the subsequent reaction was equally rapid, the primary adduct did not accumulate as much as for the AzadC/$OH^-$ reaction, which resulted in a lower absorption maximum at 248 nm. Again, the rate for the formation of the primary adduct showed a clear linear dependence on the concentration of the nucleophilic $OH^-$ ions. From the slope of the plot (Fig. S1b, right panel), a rate constant $k = 3.60 \times 10^{-4}$ ᴍ$^{-1}$ $s^{-1}$ could be determined for the attack of the $OH^-$ ions on cAzadC, which is 59 times smaller than for the analog reaction of AzadC/$OH^-$.

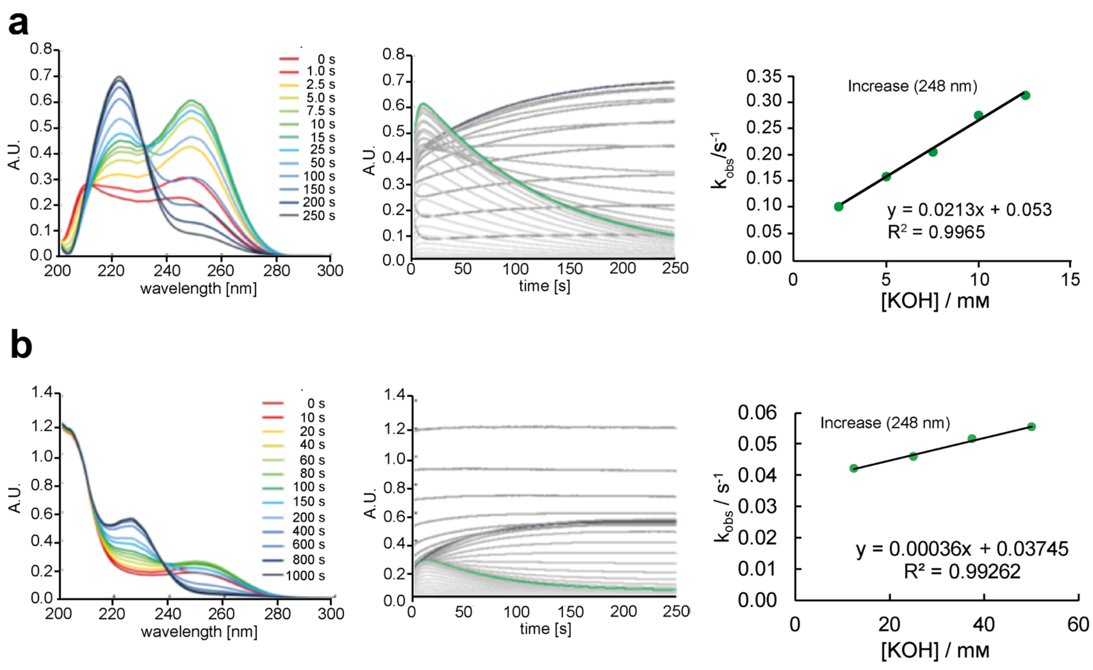


**Figure S1**. Temporal change of the UV spectra for the reactions of AzadC (a) or cAzadC (b) with $OH^-$ in aqueous solution at 20 °C and the determined rate constants $k_{obs}$ (1st order reaction) for the formation of the primary adducts. **a** [$OH^-$] = 12.5 mᴍ, [AzadC] = 0.35 mᴍ **b** [$OH^-$] = 5.0 mᴍ, [cAzadC] = 0.41 mᴍ.

Theoretical studies on AzadC **1** and cAzadC **2**

The hydrolysis of 5-aza-2'-desoxycytidine (AzadC, Decitabine, **1**) in basic aqueous solution (p*H* 11 - 12, 20 °C) proceeds approx. 59 times faster as compared to that of its carbocyclic analog cAzadC **2**. Following the currently established mechanism for aqueous degradation of **1**,[1] this may include initial hydroxide attack at the C6 position, followed by ring opening to derivative **6** (and ultimately elimination of formate). Alternatively, reversible attack of hydroxide at C4 may trigger the well-established deamination mechanism of cytosine, which eventually leads to elimination of ammonia and the C4-keto derivative **8**.[2] For **2** the same reaction pathways may be anticipated (Scheme S1).

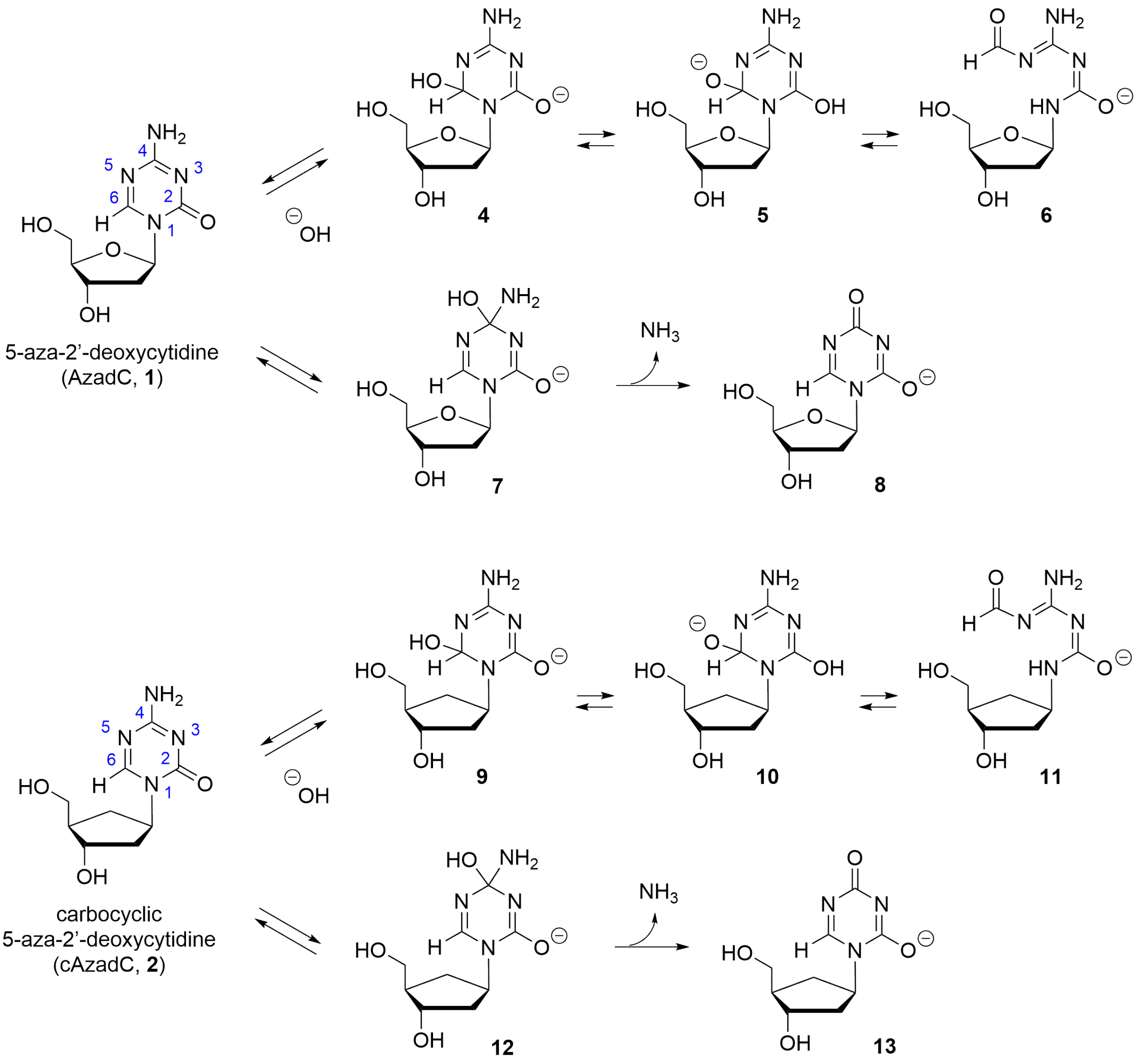


**Scheme S1**. Base hydrolysis pathways for 5-aza-2'-desoxycytidine (AzadC, Decitabine, **1**) and its carbocyclic analog **2**.

To rationalize the experimental observations, the conformational space accessible to **1** and **2** in aqueous solution has been analyzed using classical Molecular Dynamics (MD) simulations in water. For the sake of comparison, we also include 2'-deoxycytidine (dC, **3**) in these studies. MD simulations were performed using the Amber 18 software package[3] using the ff14SB and bsc1 force field parameters.[4] The parameterization of the AzadC (**1**) and cAzadC (**2**) compounds was adapted from that of dC **3**. Each molecule was inserted in a rectangular solvent box (TIP3P water type), in a total of 1026 water molecules. Each system was minimized and equilibrated for 100 ps, followed by 60 ns of production. The SHAKE algorithm[5] was employed to constrain the bond lengths involving hydrogen atoms. To control the pressure (1 atm) and the temperature (310 K) of the systems, the Berendsen barostat and the Langevin thermostat[6] were used. Periodic boundary conditions were considered. Non-bonded interaction pairs were calculated within a 10 Å. Beyond that, Coulomb interactions were treated with the Particle-Mesh Ewald (PME) method[7] and vdW interactions were truncated. The MD trajectories were analysed with the CPPTRAJ module[8], combined with the Visual Molecular Dynamics (VMD 1.9.2) program[9] for visualization and image rendering. For each MD trajectory, three different cluster analyses were carried out, in which all geometries were grouped in 10 groups, using a distance metric: the root-mean square deviation of *i)* all atoms of the molecule, *ii)* all atoms of the base ring, and *iii)* all atoms of the sugar ring. Tables S1 displays the frequency of each group as well as the dihedrals $\omega$1 (O4'-C1'-N1-C6) and $\omega$2 (O4'-C1'-C2'-C3') values of the representative geometry of each group for the AzadC, cAzadC, and dC molecules, respectively.

A clear preference for the *anti* conformation (C6-up) of the base in all molecules (ω1 > 0º) was observed. This is expected due to the fact that the DNA duplex adopts the *anti* conformation of nucleosides (well mimicked/represented by the force field). Regarding the ribose ring pucker, the C2'-endo is the most populated conformation in both dC and AzadC; however, the substitution of the O4' by a carbon changes this tendency to a C3'-endo conformation (ω2 < 0º) as shown schematically in Scheme S2.

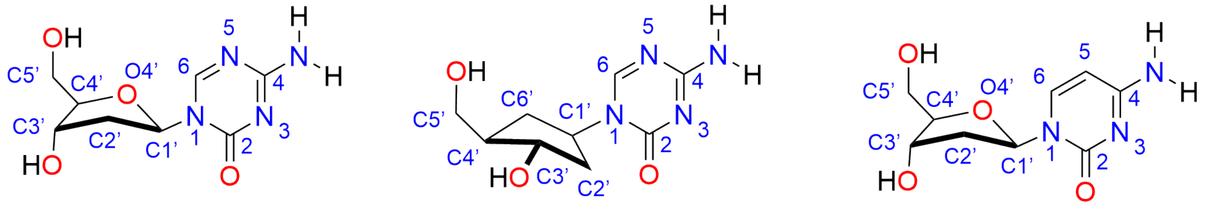

AzadC **1** cAzadC **2** dC **3**

**Scheme S2**. Schematic representations of the preferred conformations of dC (**3**), AzadC (**1**), and cAzadC (**2**) according to MD simulations in water.

**Table S1**. Frequency of each group throughout the MD simulation as well as the dihedrals ω1 (O4'-C1'-N1-C6) and ω2 (O4'-C1'-C2'-C3') of the representative geometry of each group for dC (**3**), AzadC (**1**), and cAzadC (**2**).

| cluster by rms | group | AzadC (**1**) | | | cAzadC (**2**) | | | dC (**3**) | | |
|---|---|---|---|---|---|---|---|---|---|---|
| | | frequency / % | ω1 / º | ω2 / º | frequency / % | ω1 / º | ω2 / º | frequency / % | ω1 / º | ω2 / º |
| all atoms | 1 | 81 | 41.59 | 37.02 | **42.6** | **75.8** | **-6.1** | 55.7 | 46.0 | 41.8 |
| | 2 | 7 | 73.53 | 38.41 | 29.9 | 57.7 | 42.4 | 19.6 | 25.3 | 36.8 |
| | 3 | 6 | 34.83 | 46.36 | **10.7** | **44.4** | **-30.7** | 9.6 | 59.4 | 28.6 |
| | 4 | 1.8 | 22.81 | 44.06 | 7.2 | 74.0 | 41.0 | 6.1 | 16.8 | 35.8 |
| | 5 | 1.5 | 9.02 | 13.32 | 2.5 | 54.7 | -2.9 | 5.5 | 74.4 | 40.5 |
| | 6 | 1.3 | 34.79 | -9.76 | 2.2 | 26.1 | 51.4 | 2.1 | 44.9 | -10.4 |
| | 7 | 0.5 | 85.88 | 43.35 | 1.9 | 95.3 | -26.2 | 0.7 | -120.2 | 43.6 |
| | 8 | 0.4 | 41.57 | 47.3 | 1.5 | 92.7 | 7.3 | 0.3 | 68.4 | 31.3 |
| | 9 | 0.3 | -43.56 | 35.49 | 0.7 | 76.1 | 35.2 | 0.2 | 84.0 | -10.3 |
| | 10 | 0.2 | -5.94 | -39.89 | 0.7 | 73.2 | 46.6 | 0.1 | 68.1 | 49.3 |
| base ring | 1 | 49.4 | 41.48 | 32.54 | **73.6** | **62.3** | **-6.6** | 33.3 | 46.0 | 41.76 |
| | 2 | 15.9 | 6.4 | 37.95 | **7.3** | **62.9** | **-6.5** | 23.6 | 25.34 | 36.84 |
| | 3 | 8.8 | 26.48 | 35.91 | 5.4 | 53.1 | 4.1 | 22 | 59.44 | 28.55 |
| | 4 | 7.1 | 36.88 | 44.91 | 4.7 | 77.7 | 38.5 | 0.9 | 16.78 | 35.84 |
| | 5 | 7 | 39.74 | 25.35 | 2.4 | 42.1 | -9.3 | 3.5 | 74.37 | 40.51 |
| | 6 | 4.7 | 48.67 | 39.5 | 2 | 49.2 | -4.0 | 3.2 | 44.88 | -10.35 |
| | 7 | 4 | 39.47 | 36.28 | 1.8 | 75.7 | 43.7 | 2.7 | -120.16 | 43.64 |
| | 8 | 1.6 | 44.14 | 42.99 | 1.6 | 68.4 | -22.9 | 2 | 68.39 | 31.33 |
| | 9 | 0.7 | -3.33 | 27.66 | 0.8 | 55.8 | 40.0 | 0.4 | 83.95 | -10.3 |
| | 10 | 0.7 | 27.48 | 43.66 | 0.4 | 112.0 | 49.1 | 0.2 | 68.14 | 49.33 |
| ribose ring | 1 | 33.5 | 16.08 | 34.63 | 34.2 | 51.59 | 40.51 | 48.9 | 59.61 | 41.72 |
| | 2 | 24.1 | 34.27 | 40.23 | **33** | **33.01** | **-15.28** | 23.5 | 54.85 | 44.09 |
| | 3 | 14.6 | 63.82 | 46.78 | 15 | 66 | 0.41 | 17.4 | 44.02 | 32 |
| | 4 | 8.3 | 10.25 | 24.91 | 4.1 | 54.2 | -9.66 | 3.3 | 50.45 | 33.73 |
| | 5 | 7.3 | 46.07 | 51.18 | 3.6 | 80.49 | 39.37 | 2.3 | 68.19 | 51.35 |

| | | | | | | | | | |
|---|---|---|---|---|---|---|---|---|---|
| | 6 | 5.1 | 37.74 | 28.68 | 3.5 | 3.41 | 22.89 | 2 | 78.27 | 43.25 |
| | 7 | 2.5 | 70.23 | 16.8 | 2.2 | 68.39 | -22.87 | 1.7 | 63.88 | -11.58 |
| | 8 | 2.2 | 29.29 | 30.44 | 1.6 | 55.87 | -30.08 | 0.7 | 57.99 | 16.66 |
| | 9 | 1.8 | 45.9 | 43.44 | 1.5 | 32.85 | 32.4 | 0.3 | 53.4 | 18.12 |
| | 10 | 0.5 | 42.81 | 42.43 | 1.4 | 48.1 | 49.22 | 0.1 | 46.85 | 6.81 |

Single-point quantum chemical energy calculations at the SMD(water)/B3LYP-D3/6-31+G(d,p) level of theory were then performed on the thirty representative geometries of each compound obtained in the MD simulations. Table S2 shows the total energy values in kJ mol$^{-1}$ of all geometries of the three molecules. Figure S1 illustrates the most favored geometries (lowest total energy) of AzadC (**1**) and cAzadC (**2**) solvated with all water molecules within a radius of 400 pm, while Table S3 summarizes the charge distribution of some atoms as well as the HOMO/LUMO energy data for these geometries. No significant differences can be observed in the HOMO/LUMO energies of the three molecules. Analysis of the charge distributions in the base region indicates that the substitution of the C5 by a nitrogen atom in both AzadC and cAzadC molecules increases the charge of the neighbors C4 and C6 atoms by *ca.* 0.16 and 0.27. However, no significant differences were observed in the charge of key atoms as well as in the overall charge of the base ring of both AzadC (**1**) and cAzadC (**2**): the charge of the C6 atom amounts to +0.344 in AzadC (**1**) and +0.349 in cAzadC (**2**). In addition, it was verified that the charge distribution of the key atoms is independent of the base (*anti* or *syn*) and ribose (C2'-endo or C3'-endo) conformations. Concerning the ribose group, the substitution of the O4' by a carbon on cAzadC decreased the charges of the C1' and C4' by 0.34, in comparison with both AzadC and dC molecules. As the C1' is close to C6 of the base, this particular electronegativity effect could, in combination with the conformational effect (from the ribose ring pucker change), influence or reduce the nucleophilic attack of the $HO^-$ on the C6.

**Table S2**. Single-point energy calculations at the SMD(water)/B3LYP-D3/6-31+G(d,p) level of theory. $E_{tot}$ and $\Delta E_{tot}$ values are in Hartree and kJ mol$^{-1}$ units, respectively.

| | | AzadC (1) | | cAzadC (2) | | dC (3) | |
|---|---|---|---|---|---|---|---|
| cluster by rms | group | $E_{tot}$ SMD(water)/ B3LYP-D3/ 6-31+G(d,p) | $\Delta E_{tot}$ | $E_{tot}$ SMD(water)/ B3LYP-D3/ 6-31+G(d,p) | $\Delta E_{tot}$ | $E_{tot}$ SMD(water)/ B3LYP-D3/ 6-31+G(d,p) | $\Delta E_{tot}$ |
| | 1 | -832.0798025 | 0.1 | -796.1639656 | 38.5 | -816.0173034 | 63.8 |
| | 2 | -832.0773821 | 6.4 | -796.1618055 | 44.2 | -816.0231602 | 48.5 |
| | 3 | -832.0626682 | 45.1 | -796.1780663 | 1.5 | -816.017933 | 62.2 |
| | 4 | -832.0738105 | 15.8 | -796.1652015 | 35.3 | -816.0179378 | 62.2 |
| | **5** | **-832.0798334** | **0.0** | -796.159843 | 49.4 | -816.0170988 | 64.4 |
| | 6 | -832.0670444 | 33.6 | -796.1589965 | 51.6 | -816.0232427 | 48.2 |
| | 7 | -832.0704742 | 24.6 | -796.1567612 | 57.5 | -816.0223944 | 50.5 |
| | 8 | -832.0616325 | 47.8 | -796.1693341 | 24.4 | -816.0034762 | 100.1 |
| | 9 | -832.0732924 | 17.2 | -796.1509493 | 72.7 | -816.0342452 | 19.3 |
| all atoms | 10 | -832.0634447 | 43.0 | -796.1604646 | 47.7 | -816.0062143 | 92.9 |
| | 1 | -832.0641103 | 41.3 | -796.16921 | 24.8 | -816.0132825 | 74.4 |
| | 2 | -832.0725515 | 19.1 | -796.1558364 | 59.9 | -816.0220693 | 51.3 |
| | 3 | -832.0750838 | 12.5 | -796.1743076 | 11.4 | -816.0206029 | 55.2 |
| | 4 | -832.075635 | 11.0 | -796.1464377 | 84.6 | -816.0242915 | 45.5 |
| | 5 | -832.0706781 | 24.0 | -796.1659611 | 33.3 | -816.012191 | 77.3 |
| | 6 | -832.0695186 | 27.1 | **-796.1786455** | **0.0** | -816.0254292 | 42.5 |
| | 7 | -832.0756101 | 11.1 | -796.1664473 | 32.0 | -816.0039746 | 98.8 |
| base ring | 8 | -832.0747626 | 13.3 | -796.1495968 | 76.3 | -816.022872 | 49.2 |

| | | | | | | | |
|---|---|---|---|---|---|---|---|
| | 9 | -832.0396054 | 105.6 | -796.1641989 | 37.9 | -816.0212608 | 53.4 |
| | 10 | -832.060693 | 50.3 | -796.1706543 | 21.0 | -816.0018177 | 104.5 |
| | 1 | -832.0577641 | 57.9 | -796.1677223 | 28.7 | -816.0225052 | 50.2 |
| | 2 | -832.0702711 | 25.1 | -796.1454403 | 87.2 | -816.0320508 | 25.1 |
| | 3 | -832.0656523 | 37.2 | -796.1511173 | 72.3 | **-816.0416146** | **0.0** |
| | 4 | -832.0763641 | 9.1 | -796.1695313 | 23.9 | -816.0159893 | 67.3 |
| | 5 | -832.079079 | 2.0 | -796.1779298 | 1.9 | -816.0129497 | 75.3 |
| | 6 | -832.0629524 | 44.3 | -796.1602721 | 48.2 | -816.007289 | 90.1 |
| | 7 | -832.0627538 | 44.8 | -796.1495968 | 76.3 | -816.0230578 | 48.7 |
| | 8 | -832.0744174 | 14.2 | -796.1413294 | 98.0 | -816.0189656 | 59.5 |
| | 9 | -832.0752297 | 12.1 | -796.1621835 | 43.2 | -816.0213363 | 53.2 |
| ribose ring | 10 | -832.0723323 | 19.7 | -796.1617467 | 44.4 | -816.0270276 | 38.3 |

**Table S3**. Charge distribution (Natural Bond Orbital (NBO) analysis) of some atoms as well as the HOMO/LUMO energy data at the SMD(water)/B3LYP-D3/6-31+G(d,p) level of theory, for the geometries with the lowest total energy of each compound.

| | atom | **AzadC (1)** | **cAzadC (2)** | **dC (3)** | **AzadC (1) - cAzadC (2)** | **AzadC (1) - dC (3)** | **cAzadC (2) – dC (3)** |
|---|---|---|---|---|---|---|---|
| | N1 | -0.479 | -0.446 | -0.463 | -0.03 | -0.02 | 0.02 |
| | C2 | 0.807 | 0.801 | 0.800 | 0.01 | 0.01 | 0.00 |
| | O2 | -0.731 | -0.759 | -0.755 | 0.03 | 0.02 | 0.00 |
| | N3 | -0.635 | -0.631 | -0.652 | 0.00 | 0.02 | 0.02 |
| | C4 | 0.596 | 0.589 | 0.440 | 0.01 | **0.16** | **0.15** |
| | N4 | -0.810 | -0.815 | -0.820 | 0.00 | 0.01 | 0.01 |
| | H4_1 | 0.459 | 0.456 | 0.460 | 0.00 | 0.00 | 0.00 |
| | H4_2 | 0.458 | 0.463 | 0.454 | 0.00 | 0.00 | 0.01 |
| | N5 | -0.587 | -0.592 | - | 0.01 | - | - |
| | **C6** | **0.344** | **0.349** | **0.078** | **-0.01** | **0.27** | **0.27** |
| | H6 | 0.267 | 0.280 | 0.279 | -0.01 | -0.01 | 0.00 |
| base | total | -0.311 | -0.306 | -0.179 | -0.01 | -0.13 | -0.13 |
| | C1' | 0.255 | -0.083 | 0.259 | **0.34** | 0.00 | **-0.34** |
| | H1' | 0.275 | 0.307 | 0.264 | -0.03 | 0.01 | 0.04 |
| | C2' | -0.527 | -0.514 | -0.529 | -0.01 | 0.00 | 0.02 |
| | C3' | 0.051 | 0.078 | 0.038 | -0.03 | 0.01 | 0.04 |
| | O3' | -0.830 | -0.833 | -0.828 | 0.00 | 0.00 | 0.00 |
| | C4' | 0.029 | -0.308 | 0.039 | **0.34** | -0.01 | **-0.35** |
| | H4' | 0.275 | 0.274 | 0.266 | 0.00 | 0.01 | 0.01 |
| | C5' | -0.140 | -0.117 | -0.131 | -0.02 | -0.01 | 0.01 |
| ribose | O5' | -0.824 | -0.846 | -0.823 | 0.02 | 0.00 | -0.02 |
| | base | *anti* / C6-up | *anti* / C6-up | *anti* / C6-up | | | |
| conformation | ribose | C2'-endo | C3'-endo | C2'-endo | | | |

| | | | | | | | |
|---|---|---|---|---|---|---|---|
| MO | HOMO | -0.26307 | -0.26085 | -0.24356 | 0.00 | -0.02 | -0.02 |
| | LUMO | -0.04119 | -0.06335 | -0.04255 | 0.02 | 0.00 | -0.02 |
| dihedral | ω1 | 9.057 | 49.215 | 44.081 | | | |
| | ω2 | 13.267 | -4.051 | 31.959 | | | |

All thirty representative geometries of each compound were then optimized at the SMD(water)/B3LYP-D3/6-31+G(d,p) level of theory. Figure S2 illustrates the thermo-dynamically most favored geometries of each compound, whilst Table S4 summarizes the charge distribution of some atoms as well as the HOMO/LUMO energy data for these geometries. The results are quite similar to those obtained in the single-point energy calculations. Again, it can be seen that AzadC (**1**) and cAzadC (**2**) share a rather similar charge distribution in the base, which also leads to comparable electrostatic potential energy maps for both. From this type of ground state data it is thus not immediately possible to derive significantly different reactivities towards nucleophiles at the C6 position of the base.

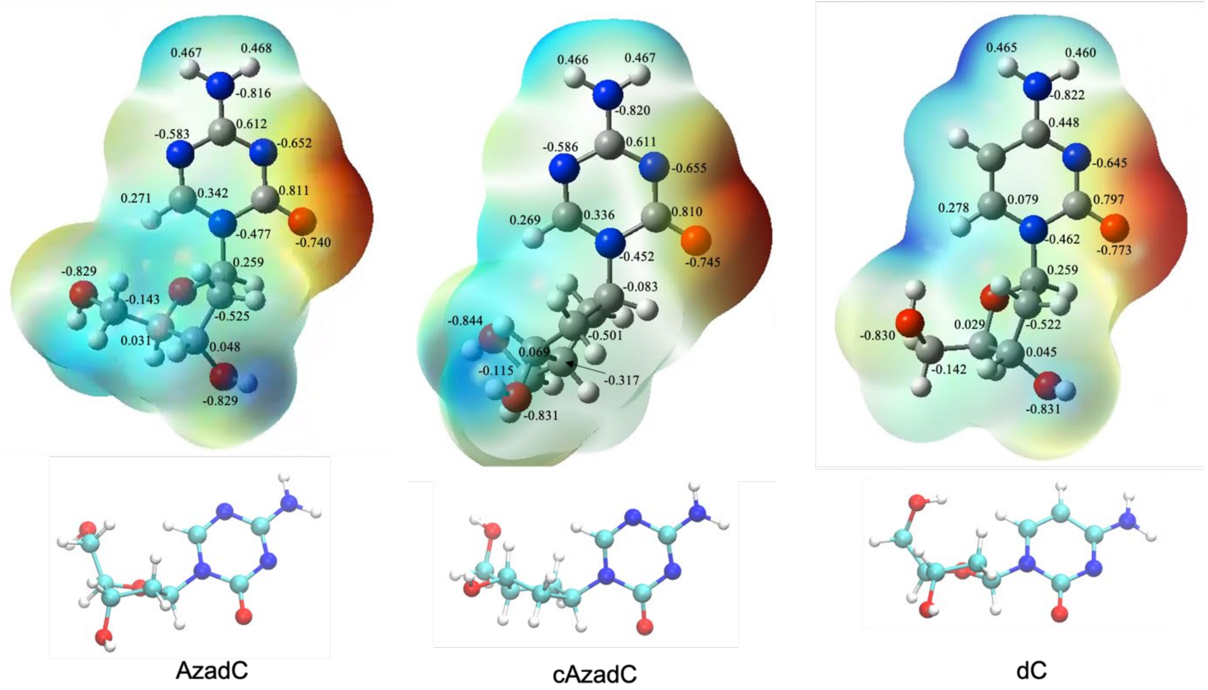


**Figure S2**. Thermodynamically most favored geometries of AzadC (**1**), cAzadC (**2**), and dC (**3**) obtained at the SMD(water)/B3LYP-D3/6-31+G(d,p) level of theory. The electrostatic potential energy (ESP) map and the charges (NBO analysis) of the key atoms are also shown.

**Table S4**. Charge distribution of some atoms as well as the HOMO/LUMO energy data for the thermodynamically most favored geometries of each compound.

| | atom | **AzadC (1)** | **cAzadC (2)** | **dC (3)** | **AzadC (1) - cAzadC (2)** | **AzadC (1) - dC (3)** | **cAzadC (2) – dC (3)** |
|---|---|---|---|---|---|---|---|
| base | N1 | -0.477 | -0.452 | -0.462 | -0.03 | -0.01 | 0.01 |
| | C2 | 0.811 | 0.810 | 0.797 | 0.00 | 0.01 | 0.01 |
| | O2 | -0.740 | -0.745 | -0.773 | 0.00 | 0.03 | 0.03 |
| | N3 | -0.652 | -0.655 | -0.645 | 0.00 | -0.01 | -0.01 |
| | C4 | 0.612 | 0.611 | 0.448 | 0.00 | 0.16 | 0.16 |
| | N4 | -0.816 | -0.820 | -0.822 | 0.00 | 0.01 | 0.00 |
| | H4_1 | 0.467 | 0.466 | 0.465 | 0.00 | 0.00 | 0.00 |
| | H4_2 | 0.468 | 0.467 | 0.460 | 0.00 | 0.01 | 0.01 |
| | N5 | -0.583 | -0.586 | - | 0.00 | - | - |

| | | | | | | | |
|---|---|---|---|---|---|---|---|
| | **C6** | **0.342** | **0.336** | **0.079** | **0.01** | **0.26** | **0.26** |
| | H6 | 0.271 | 0.269 | 0.278 | 0.00 | -0.01 | -0.01 |
| | total | -0.29845 | -0.298 | -0.175 | 0.00 | -0.12 | -0.12 |
| | C1' | 0.259 | -0.083 | 0.259 | 0.34 | 0.00 | -0.34 |
| | H1' | 0.275 | 0.297 | 0.270 | -0.02 | 0.01 | 0.03 |
| | C2' | -0.525 | -0.501 | -0.522 | -0.02 | 0.00 | 0.02 |
| | C3' | 0.048 | 0.069 | 0.045 | -0.02 | 0.00 | 0.02 |
| | O3' | -0.829 | -0.831 | -0.831 | 0.00 | 0.00 | 0.00 |
| | C4' | 0.031 | -0.317 | 0.029 | 0.35 | 0.00 | -0.35 |
| | H4' | 0.265 | 0.272 | 0.264 | -0.01 | 0.00 | 0.01 |
| | C5' | -0.143 | -0.115 | -0.142 | -0.03 | 0.00 | 0.03 |
| ribose | O5' | -0.829 | -0.844 | -0.830 | 0.02 | 0.00 | -0.01 |
| | base | *anti* / C6-up | *anti* / C6-up | *anti* / C6-up | | | |
| conformation | ribose | C2'-endo | C3'-endo | C2'-endo | | | |
| | HOMO | -0.26345 | -0.26595 | -0.24254 | 0.00 | -0.02 | -0.02 |
| MO | LUMO | -0.05163 | -0.05008 | -0.04296 | 0.00 | -0.01 | -0.01 |
| | ω1 | 38.632 | 68.251 | 40.259 | | | |
| dihedral | ω2 | 29.115 | -5.087 | 39.299 | | | |

Preparation of AzadC **1** and cAzadC **2** for cell culture experiments

AzadC **1** was purchased (Carbosynth NA02969) and used without further purification. cAzadC 2 was synthesized as previously described by *Wildenhof et al., 2019*[10]. Both compounds were dissolved in DMSO to a final concentration of 10 mM. Aliquots (10 µL) were made and stored at -80 °C. Directly before usage, the required volume of 10 mM stock solution was thawed and diluted to 100 µM with Milli-Q $H_2O$.

Cell culture MOLM-13

MOLM-13 cells (Leibniz Institute DSMZ-German Collection of Microorganisms and Cell Cultures; AML cell line[11]) were cultivated at 37°C in water saturated, $CO_2$-enriched (5%) atmosphere. RPMI 1640 (Sigma-Aldrich R0883), containing 20% (v/v) fetal bovine serum (FBS) (Invitrogen 10500-064) and 1% (v/v) L-alanyl-L-glutamine (Sigma-Aldrich G8541) were used as growing medium. When reaching a density of $2x10^6$ cells/mL, the cells were routinely passaged to a density of 0.25 – 0.4 x $10^6$ cells/mL. Cells were tested at least once in two months for Mycoplasma contamination using Mycoplasma Detection Kit (Jena Bioscience PP-401L).

Treatment of MOLM-13 with AzadC **1** or cAzadC **2**

For all experiments, cells were seeded at a concentration of 0.5 x $10^6$ cells/mL and directly treated with AzadC **1** or cAzadC **2** using the indicated concentration and incubation time during which the medium was not renewed. Cells that were treated with 0.03% (v/v) of DMSO (final concentration in the medium), which is the DMSO dose the cells were exposed to when treated with 3 µM of AzadC 1 or cAzadC 2, served as a negative control in all experiments that involved MOLM-13.

Isolation of nuclear proteins

For the preparation of nuclear extracts 2.5 x $10^6$ cells were seeded and treated with 0.5 µM of AzadC **1** or cAzadC **2** or 0.03% of DMSO for 40 h. After treatment, the cells were harvested and nuclear extracts were prepared as previously described by *Dignam et al., 1983*[12] with the modification that every buffer was supplemented with Phosphatase Inhibitor Cocktail 2 (Sigma-Aldrich P5726) and Phosphatase Inhibitor Cocktail 3 (Sigma-Aldrich P0044), 1:100 each. Afterwards, the protein concentration was determined using a Bradford assay (Biorad #5000006) as described by the manufacturer. SDS loading buffer (final concentration 50 mM Tris pH 6.8, 100 mM DTT, 2% (w/v) SDS, 10% (v/v) glycerol, 0.1% (w/v) bromophenol blue) was added and the samples were incubated for 5 min at 92 °C before being stored at -20 °C. Before loading the samples on a polyacrylamide gel, the samples were heated for additional 2 min at 92 °C and vortexed thoroughly.

Western blotting

15 µg of nuclear extract in SDS loading buffer were loaded on a 4-15% precast polyacrylamide gel (Bio-Rad #4561083EDU) and Color-coded Prestained Protein Marker, Broad Range (10-250 kDa) (New England Biolabs P7719S) was used as a protein standard. The gel was run at constant 150 V for 60 min in SDS running buffer (25 mM Tris, 192 mM glycine, 0.1% (w/v) SDS). For blotting, we used a PVDF blotting membrane (GE Healthcare Amersham Hybond P0.45 PVDG membrane 10600023) and pre-cooled Towbin blotting buffer (25 mM Tris, 192 mM glycine, 20% (v/v) methanol, 0.038% (w/v) SDS). The membrane was activated for 1 min in methanol, washed with Milli-Q water and equilibrated for additional 1 min in Towbin blotting buffer; the Whatman gel blotting papers (Sigma-Aldrich WHA 10426981) were equilibrated for 15 min in Towbin buffer and the precast gel was equilibrated for 5 min in Towbin buffer after the run. Western blotting (tank (wet) electro transfer) was performed at 4°C for 9 h at constant 35 V. After blotting, the PVDF membrane was blocked for 0.5 - 1 h at room temperature using 5% (w/v) milk powder in TBS-T (20 mM Tris pH = 7.5, 150 mM NaCl, 0.1% (v/v) Tween-20). The primary antibodies were diluted in 5 mL of 5% (w/v) milk powder in TBS-T. The blocking suspension was discarded, and the diluted primary antibodies were added for 12 h at 4°C and shaking. After incubation, the primary antibodies were discarded, and the membrane was washed three times ten minutes with TBS-T. HRP-conjugated secondary antibodies were diluted in 5% (w/v) milk powder in TBS-T and added for 1 h at room temperature under shaking. Afterwards, the membrane was washed two times with TBS-T and one time with TBS (TBS-T without Tween-20) before SuperSignal West Pico Chemiluminescent Substrate (Thermo Scientific 34077) was used for imaging. Western blots were imaged using Amersham Imager 680 (auto exposure mode).

For imaging the same blot multiple times using different antibodies, the membrane was directly stripped after imaging. To this end, the membrane was put in TBS-T and the buffer was heated in a microwave until boiling. Afterwards, the buffer was discarded and the procedure was repeated in total three times. After stripping, the membrane was blocked again using 5% (w/v) milk powder in TBS-T and the protocol followed the above described procedure.

Primary antibodies

- Anti-DNMT1 antibody, Active Motif 61457 clone 2C1, rat monoclonal antibody, 1:1000
- Anti-phospho-Histone-H2AX (γH2AX, Ser-139) antibody, Millipore 05-636-1 clone 7BW301, mouse monoclonal antibody, 1:1000
- Anti-Histone-H3 antibody, Cell Signaling Technology 4499S clone D1H2, rabbit monoclonal antibody, 1:1000

Secondary antibodies

- HRP-conjugated anti-mouse IgG, Sigma-Aldrich AP130P, 1:5000
- HRP-conjugated anti-rabbit IgG, Sigma-Aldrich A0545, 1:5000
- HRP-conjugated anti-rat IgG, Cell Signaling technology 7077S, 1:5000

Isolation of gDNA and UHPLC-QQQ-MS

For the determination of mdC levels, 1.5 x $10^6$ cells were seeded and treated with either 0.5 µM, 1.0 µM or 3.0 µM of AzadC **1** or cAzadC **2** or with only 0.03% of DMSO for 72 h. After 72 h of incubation time, the cells were harvested, gDNA was isolated and UHPLC-QQQ-MS measurements were performed as previously described in *Traube et al., 2019*.[13] Three biological independent experiments were performed for each experimental condition.

MTT assay

For the MTT assay, 5 x $10^4$ cells were seeded in 100 µL RPMI/20% FBS medium that did not contain phenol red. The assay was performed as described previously[14]. Each timepoint (24 h, 48 h or 72 h) included samples for 0.03% DMSO-treated cells and cells that were treated with 0.5 µM, 1.0 µM or 3.0 µM of either AzadC **1** or cAzadC **2**. Each timepoint was measured individually and after measuring the absorption at 570 nm, the average of the technical replicates was calculated and the absorption of the AzadC **1** or cAzadC **2** treated cells was set in relation to the absorption measured for the DMSO-only treated cells, resulting in the relative metabolic activity. Three biologically independent experiments were performed per time point and each sample was measured in technical quadruplicates.

Alkaline and neutral comet assay

DNA single-strand breaks induced by AzadC 1 and cAzadC 2 were detected and quantified via single-cell gel electrophoresis under alkaline conditions, as described.[15] Shortly, MOLM13 cells were exposed to 0.5 µM cAzadC and 0.5 µM AzadC and 60 h later the comet assay was performed. As positive control, cells were exposed to 150 µM tert-Butyl hydroperoxide (*t*-BuOOH) for 1 h. Cells were imbedded in 0.5 % low melting point agarose in PBS and spread onto glass slides coated with 1.2 % agarose. All following steps were performed at 4 °C. Cells were lysed in lysing solution (2.5 M NaCl, 100 mM EDTA, 10 mM Tris, 1% Na-Laurylsarcosinate, 1% Tryton X-100) at a pH of 10 for the alkaline comet and at a pH of 7.5 for the neutral comet. For the alkaline comet, DNA was unwound for 25 min and electrophoresed for 15 min in the same solution (0.3 N NaOH, 6 mM EDTA, pH>13) at 0.74 V/cm and 300 mA. Slides were then neutralized (0.4 M Tris, pH 7.5), fixed in 100 % methanol and dried. For the neutral comet, slides were electrophoresed for 25 min in electrophoresis buffer (90 mM Tris, 90 mM boric acid,2 mM EDTA, pH= 7.5), fixed in 100% methanol and then dried. All following steps were performed at room temperature. DNA was stained using propidium iodide (50 µg/ml) and 50 cells per slide were evaluated using a fluorescence microscope and the Comet IV software (Perceptive Imaging, Liverpool, UK). Data are expressed as tail intensity or tail moment, which denotes the percentage of DNA in the tail multiplied by the length between the centre of the head and tail.[16]

Statistical Analysis

Statistical analysis was performed using GraphPad Prism 9.

### Additional supplementary figures

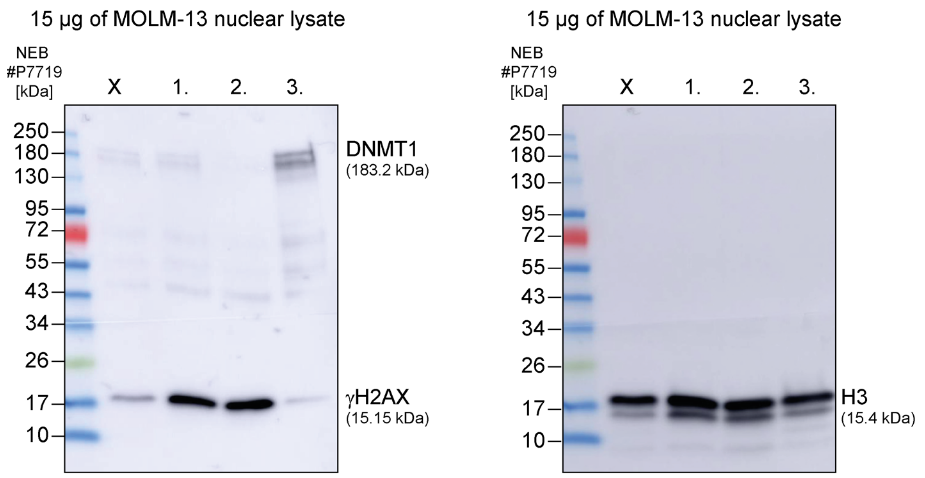


1. 0.5 µM AzadC **1** (40 h)
2. 0.5 µM cAzadC **2** (40 h)
3. 0.03% DMSO treated MOLM-13 (ctrl.)

1. 0.5 µM AzadC **1** (40 h)
2. 0.5 µM cAzadC **2** (40 h)
3. 0.03% DMSO treated MOLM-13 (ctrl.)

**Figure S3.** Depicted are the entire images for the western blots in Fig. 3 and 4. X represents a treatment that is not included in this study.

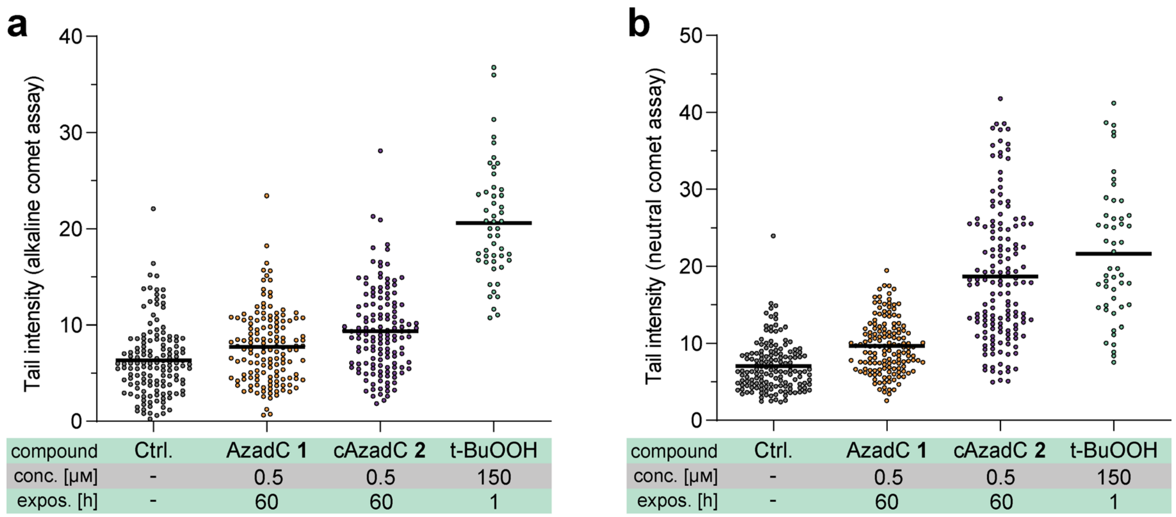


**Figure S4.** Alkaline comet assay (DNA single strand breaks, a) and neutral comet assay (DNA double strand breaks, b) including the t-BuOOOH positive control. Each dot represents the tail intensity measurement of one individual cell. Measurements from three independent experiments were analyzed and are displayed in that graph.

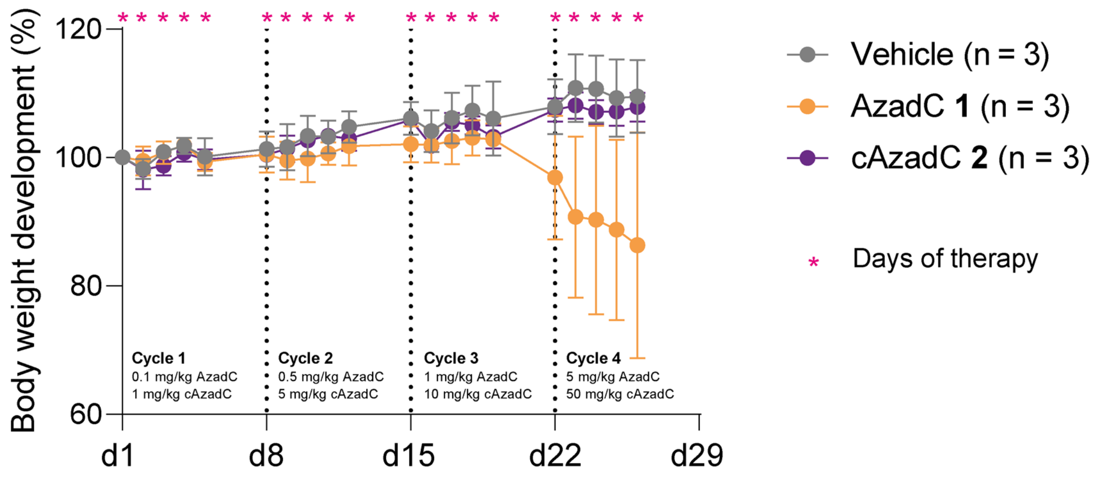


**Figure S5.** Bodyweight development of mice, which are treated with AzadC **1** or cAzadC **2** in different concentrations. Each treatment cycle consists of treatment with the compound for five consecutive days, followed by a treatment break for two days. The bodyweight of each mice was normalized to the body weight the mouse had at the begin of therapy (d1).

**References Supplement**


[1] D. K. Rogstad, J. L. Herring, J. A. Theruvathu, A. Burdzy, C. C. Perry, J. W. Neidigh, L. C. Sowers, *Chem. Res. Toxicol.* **2009**, *22*, 1194-1204.

[2] C. S. Nabel, S. A. Manning, R. M. Kohli, *ACS Chem. Biol.* **2012**, *7*, 20-30.

[3] D. A. Case, R. C. Walker, T. E. Cheatham III, C. Simmerling, A. Roitberg, K. M. Merz, R. Luo, T. Darden, J. Wang, R. E. Duke, D. R. Roe, S. LeGrand, J. Swails, A. W. Götz, J. Smith, D. Cerutti, S. R. Brozell, T. Luchko, V. W. D. Cruzeiro, D. Ghoreishi, G. Monard, C. Sagui, F. Pan, G. A. Cisneros, Y. Miao, J. Shen, R. Harris, Y. Huang, C. Lin, D. J. Mermelstein, P. Li, A. Onufriev, S. Izadi, R. M. Wolf, X. Wu, H. Gohlke, S. Schott-Verdugo, N. Homeyer, R. Qi, L. Xiao, H. Wei, D. A. Greene, T. Lee, D. York, J. Liu, H. Nguyen, I. Omelyan, A. Kovalenko, M. Gilson, I. Ben-Shalom, C. Nguyen, R. Salomon-Ferrer, T. Kurtzmann, P. A. Kollman, University of California, San Francisco.
